# Chromosome (mis)segregation is biased by kinetochore size

**DOI:** 10.1101/278572

**Authors:** Danica Drpic, Ana Almeida, Paulo Aguiar, Helder Maiato

## Abstract

**Summary:**

Aneuploidy, the gain or loss of chromosomes, arises through problems in chromosome segregation during mitosis or meiosis and has been implicated in cancer and developmental abnormalities in humans [1]. Possible routes to aneuploidy include a compromised spindle assembly checkpoint (SAC), cohesion defects, centrosome amplification, as well as improper kinetochore-microtubule attachments [2]. However, none of these established routes takes into account the intrinsic features of the kinetochore - the critical chromosomal interface with spindle microtubules. Kinetochore size and respective microtubule binding capacity varies between different animal and plant species [3-10], among different chromosomes from the same species (including humans) [3, 11-15], and in response to microtubule attachments throughout mitosis [16-18]. How kinetochore size impacts chromosome segregation remains unknown. Addressing this fundamental question in human cells is virtually impossible, because most of the 23 pairs of chromosomes cannot be morphologically distinguished, while detection of 2-3 fold differences in kinetochore size is limited by diffraction and cannot be resolved by conventional light microscopy in living cells. Here we used the unique cytological attributes of female Indian muntjac, the mammal with the lowest known chromosome number (n=3), to track individual chromosomes with distinct kinetochore sizes throughout mitosis. We found that chromosomes with larger kinetochores bi-orient more easily and are biased to congress to the equator in a motor-independent manner. However, they are also more prone to establish erroneous merotelic attachments and lag behind during anaphase. Thus, we uncovered an intrinsic kinetochore feature - size - as an important determinant of chromosome segregation fidelity.

## Results and Discussion

Kinetochore size is generally determined by the length of α-satellite DNA, the presence of a CENP-B-box, and the proportional incorporation of CENP-A at centromeres [19-22]. Additional size changes are due to an expandable module formed by CENP-C and outer kinetochore proteins involved in SAC signaling, as well as motor proteins, such as CENP-E and cytoplasmic dynein [12, 17,23]. In humans, the length of α-satellite DNA arrays at centromeres ranges from 200 kb on the Y chromosome, to >5 Mb on chromosome 18 [24], leading to an ∼3 fold variability in the amount of centromeric CENP-A and respective kinetochore size among different chromosomes [11-14, 19-21]. To investigate the relevance of kinetochore size for chromosome congression and segregation during mitosis we took advantage from the unique cytological features of the Indian muntjac (IM), a small deer whose females have the lowest known chromosome number (n=3) in mammals [25]. As the result of tandem fusions during evolution, IM chromosomes are large and morphologically distinct (one metacentric and two acrocentric pairs). Specifically relevant for our purposes, one of the acrocentric chromosomes resulting from the fusion between chromosomes 3 and X contains an unusually large compound kinetochore that can bind more than 100 microtubules [5, 25,26] (Fig. 1A). Like in humans, kinetochore size among the different IM chromosomes scaled with the amount of CENP-A and Ndc80/Hec1 and varied 2-3 fold (Fig. 1B,C). To track chromosomes and kinetochores in living cells we established a culture of hTERT-immortalized primary fibroblasts from a female IM [27] stably expressing histone H2B-GFP or CENP-A-GFP, respectively. Spindle microtubules were visualized with 20 nM SiR-tubulin [28], which did not interfere with normal mitotic progression in this system (Figs. 1D, E and 2A, movies S1 and S2).

**Fig. 1.**
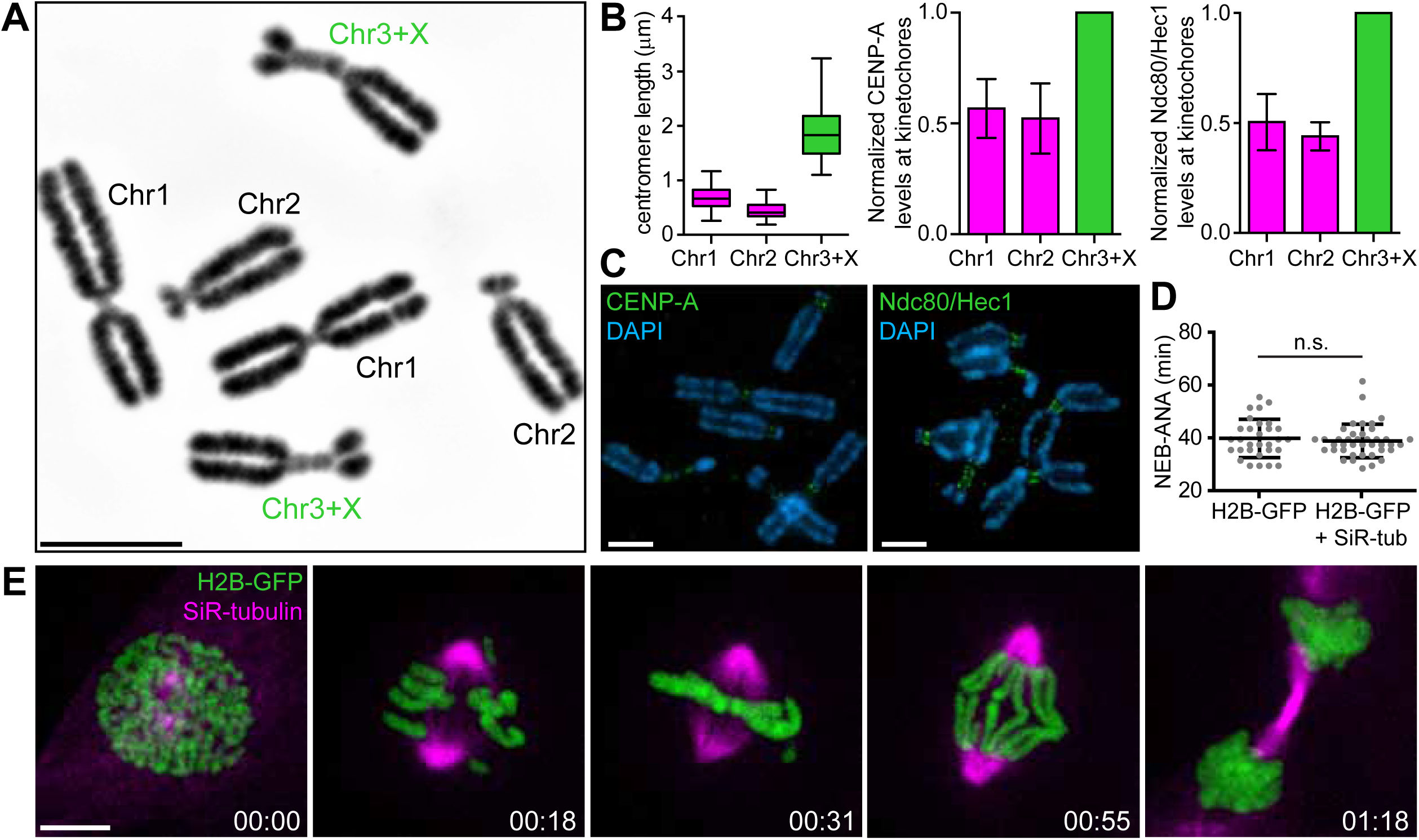
Chromosome and kinetochore size variability in Indian muntjac cells. (**A**) Normal karyotype of female Indian muntjac (IM) fibroblasts. Scale bar = 5 μm. (**B**) Linear centromere length of IM chromosomes, measured from chromosome spreads (box-whisker plots, left) and respective quantification of CENP-A (middle) and Ndc80/Hec1 (right) levels at IM kinetochores, relative to chromosome 3+X. (**C**) Immunofluorescence of IM chromosome spreads (blue) to detect CENP-A or Ndc80/Hec1 (green). Scale bars = 5 μm. (**D**) Mitotic progression from nuclear envelope breakdown (NEB) until anaphase onset (ANA) in IM fibroblasts is unperturbed by treatment with 20 nM SiR-tubulin. (**E**) Live-cell imaging of IM fibroblasts stably expressing H2B-GFP (green) and treated with 20 nM SiR-tubulin dye (magenta). Scale bar = 5 μm. Time = h:min.

In mammals, the initial contacts between mitotic spindle microtubules and kinetochores take place during prometaphase of mitosis after nuclear envelope breakdown (NEB). Scattered chromosomes then align at the spindle equator by a process known as chromosome congression [29]. When chromosomes are favorably positioned between the spindle poles they establish end-on kinetochore-microtubule attachments and congress after bi-orientation. In contrast, more peripheral chromosomes are laterally transported along spindle microtubules by the plus-end-directed kinetochore motor CENP-E (kinesin-7) [30, 31]. To directly test the implications of kinetochore size for chromosome congression we treated IM fibroblasts for 1 h with the CENP-E inhibitor GSK923295 [32] and followed their entry and progression throughout mitosis (Fig. 2B and movie S3). We found three different scenarios: 1) very few cells whose centrosomes did not fully separate before NEB showed all chromosomes at the poles (2/28 cells, 6 independent experiments), in line with our previous observations in human cells [31]; 2) some cells aligned all their chromosomes at the metaphase plate soon after NEB (6/28 cells, 6 independent experiments); and 3) most cells showed at least one chromosome, either with a small or large kinetochore, which remained at the poles (20/28 cells, 6 independent experiments). These data indicate that any chromosome is competent to use either the CENP-E-dependent or -independent pathway to congress, regardless of kinetochore size.

**Fig. 2.**
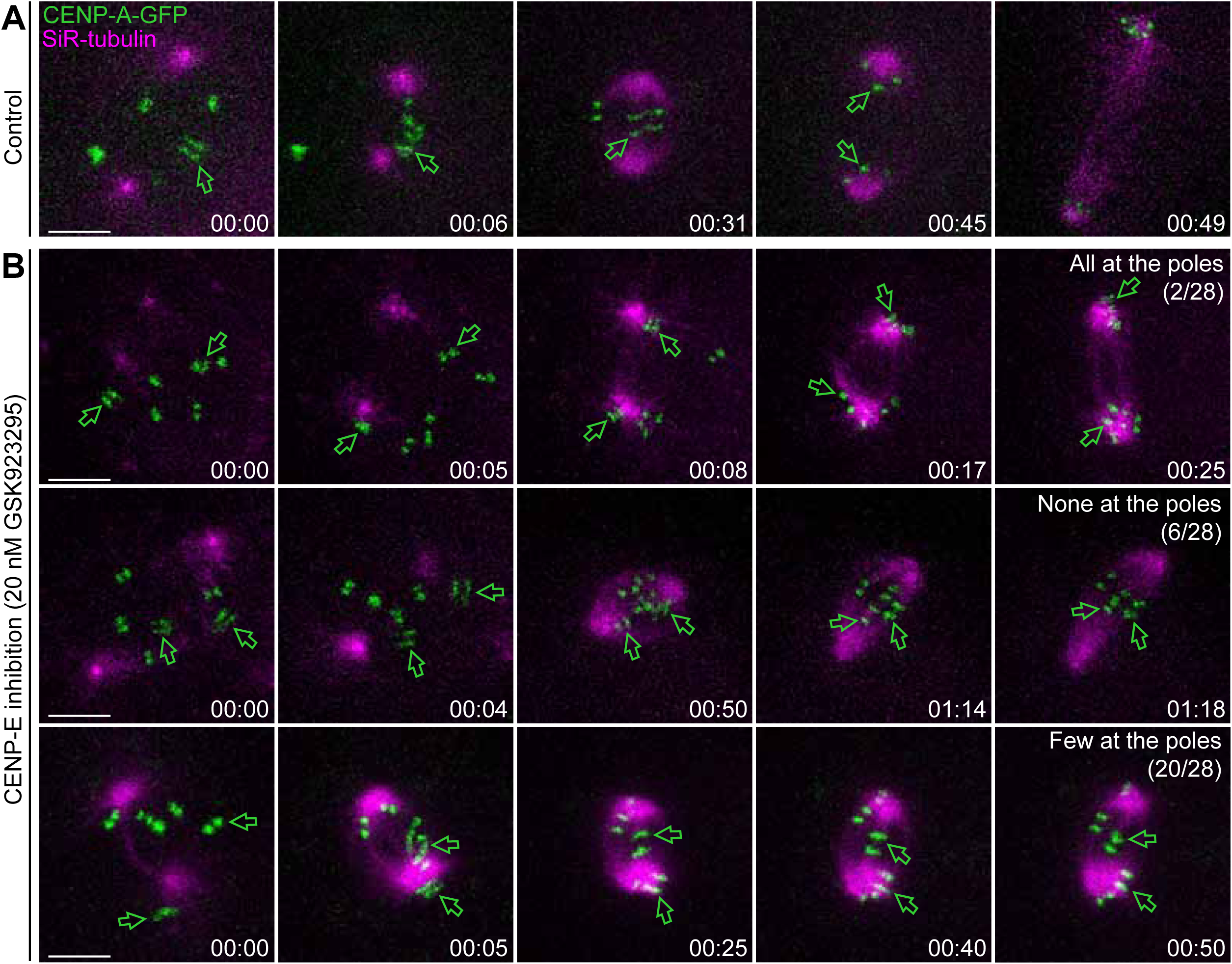
Any chromosome can use either the CENP-E-dependent or -independent pathway to congress, regardless of kinetochore size. (**A**) Control IM fibroblasts stably expressing CENP-A-GFP (green) and treated with 20 nM SiR-tubulin (magenta). Scale bar = 5 μm. Time = h:min. (**B**) IM fibroblasts stably expressing CENP-A-GFP (green) and treated with 20 nM SiR-tubulin (magenta) after CENP-E inhibition with 20 nM GSK923295. Green arrowheads indicate chromosomes with large kinetochores. Scale bars = 5 μm. Time = h:min.

Next, we determined the number of chromosomes with small or large kinetochores at the pole after CENP-E inhibition by immunofluorescence in fixed cells containing only 6 chromosomes (Fig. 3A). Based on these data we constructed a joint probability table involving two defined random variables representing the number of chromosomes with small and large kinetochores found at the pole after CENP-E inhibition (table S1). We concluded that: 1) the number of chromosomes with small and large kinetochores at the pole could be described by a binomial distribution (Fig. 3B), meaning that the fate of each individual chromosome was independent of the other chromosomes of the same class (i.e. with small or large kinetochores); 2) the state of the chromosomes with small kinetochores did not influence the state of the chromosomes with large kinetochores and vice-versa; and 3) the probability of each individual chromosome with small kinetochores to stay at the pole was approximately twice the probability of a chromosome with a large kinetochore (regardless of the number of chromosomes with small or large kinetochores): 0.19 ± 0.034 vs. 0.11 ± 0.048, respectively (mean ± s.d., n=621 cells, 6 independent experiments, p=0.0067, t test; Fig. 3C).

**Fig. 3.**
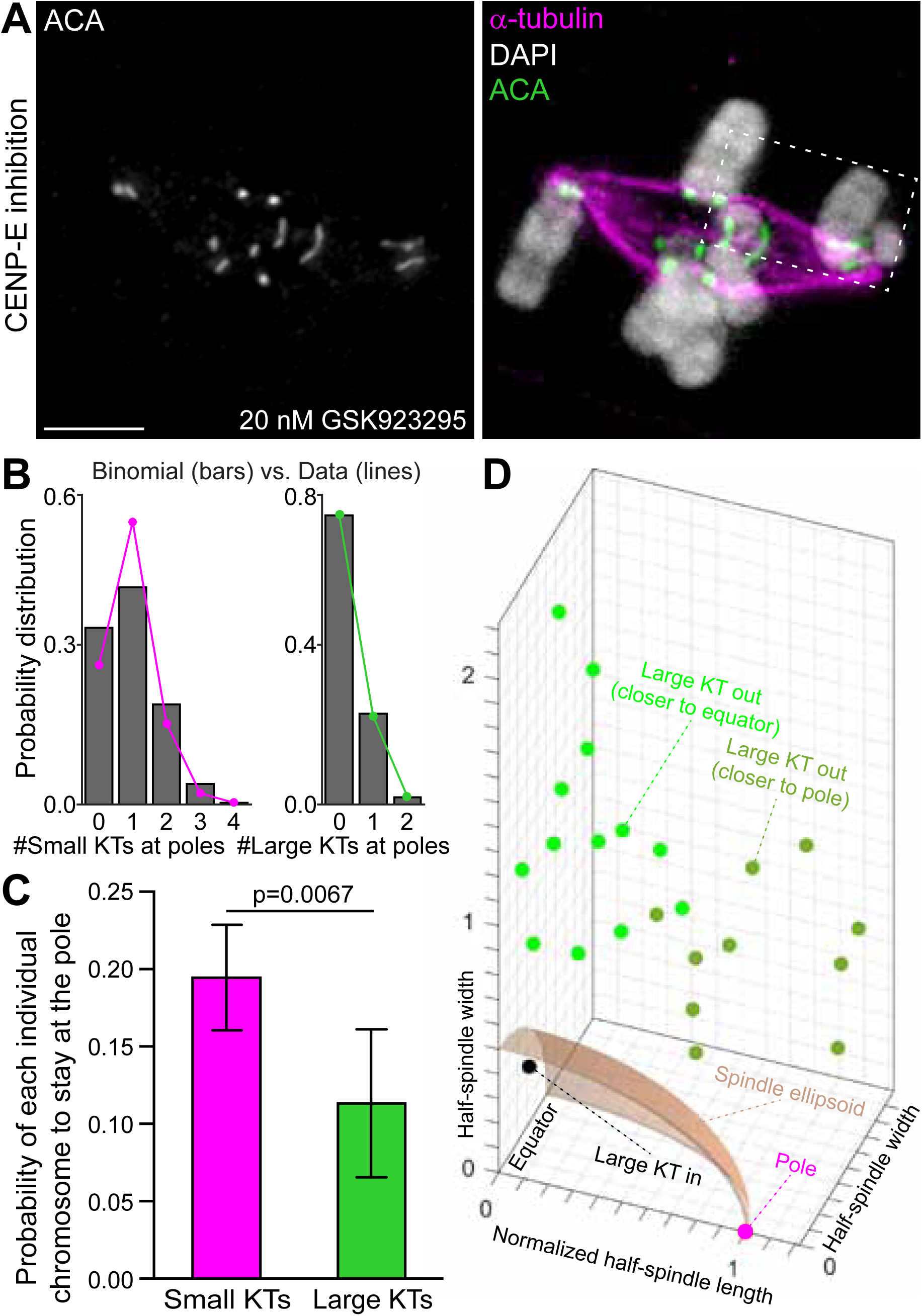
Chromosomes with a larger kinetochore rely less on CENP-E and are biased to congress after bi-orientation. (**A**) Immunofluorescence of an IM fibroblast after CENP-E inhibition showing chromosomes (DAPI, white in merged image), kinetochores (ACA, white; green in merged image) and microtubules (α-tubulin, magenta in merged image). Scale bar = 5 μm. (**B**) Quantification of the number of chromosomes with small or large kinetochores (KTs) at the pole after CENP-E inhibition by immunofluorescence in fixed cells with 6 chromosomes only (magenta and green lines) and respective theoretical prediction based on a binomial distribution (grey bars). (**C**) Probability of each individual chromosome with small or large kinetochores (KTs) to stay at the pole upon CENP-E inhibition. (**D**) Location of chromosomes with large kinetochores (KTs) after CENP-E inhibition obtained from 4D (x, y, z, t) tracking, to determine their position relative to the poles at NEB and the forming mitotic spindle (see dashed box in A for reference). Scale bar = 5 μm.

In human cells, >96% of the chromosomes relying on CENP-E for congression are normally excluded from the spindle region and locate near one of the spindle poles at NEB [31]. To rule out that the observed bias for IM chromosomes with large kinetochores to align independently of CENP-E was due to a tendency to localize between the two poles at NEB, we performed four-dimensional (4D, x,y,z,t) tracking of chromosomes with large kinetochores after CENP-E inhibition in living cells (n = 23 large kinetochore pairs, 13 cells). We found that 22/23 IM chromosomes with large kinetochores were excluded from the spindle region and were nearly randomly positioned along the spindle axis at NEB (45% of the kinetochores were closer to the poles vs. 55% of the kinetochores that were closer to the spindle equator; Fig. 3D and movie S4). Overall, these data indicate that chromosomes with a larger kinetochore rely less on CENP-E motor activity and are biased to congress after bi-orientation, despite being randomly located at the spindle periphery at NEB.

Recent computational modelling predicted that changes in kinetochore size in response to microtubule attachments affect both error formation and the efficiency of spindle assembly in human cells [18]. To directly test this prediction and investigate whether chromosomes with large kinetochores are more prone to establish erroneous attachments with spindle microtubules we set up a monastrol treatment/washout assay [33] in IM fibroblasts to inhibit kinesin-5 and spindle bipolarity, thereby preventing the formation of amphitelic attachments (Fig. 4A, B and movie S5). After monastrol washout, cells were released into fresh medium with or without the Aurora B inhibitor ZM447439 (and the proteasome inhibitor MG132 to prevent anaphase onset), to partially inhibit error correction without interfering with bipolar spindle assembly (fig. S1A-C), and subsequently fixed. For each cell treated with ZM447439 and MG132, we calculated the fraction of each chromosome group (with small or large kinetochores) with merotelic (microtubules from both poles attached to the same kinetochore) or syntelic (microtubules from the same pole attached to both sister kinetochores) attachments (Fig. 4C). We found an equivalent low frequency of syntelic attachments for chromosomes with small or large kinetochores (1.9% vs. 1.4%, respectively; n=207 cells, pool of 3 independent experiments). However, the frequency of chromosomes with large kinetochores that established erroneous merotelic attachments was several fold higher when compared with chromosomes with small kinetochores (7.0% vs. 1.6%, respectively). These results could be explained either by differences in the formation or in the correction of erroneous merotelic attachments that could scale with kinetochore size. To distinguish between these possibilities, we measured Aurora B activity in chromosomes with different kinetochore sizes by quantifying the levels of phosphorylated Knl1, a bona fide Aurora B substrate at the kinetochores [34]. We were unable to detect any significant difference in Aurora B activity between chromosomes with small or large kinetochores (n=76 small KTs, n=63 large KTs, 50 cells, Mann-Whitney test) (Fig. 4D). We concluded that chromosomes with larger kinetochores are more prone to establish erroneous merotelic attachments.

**Fig. 4.**
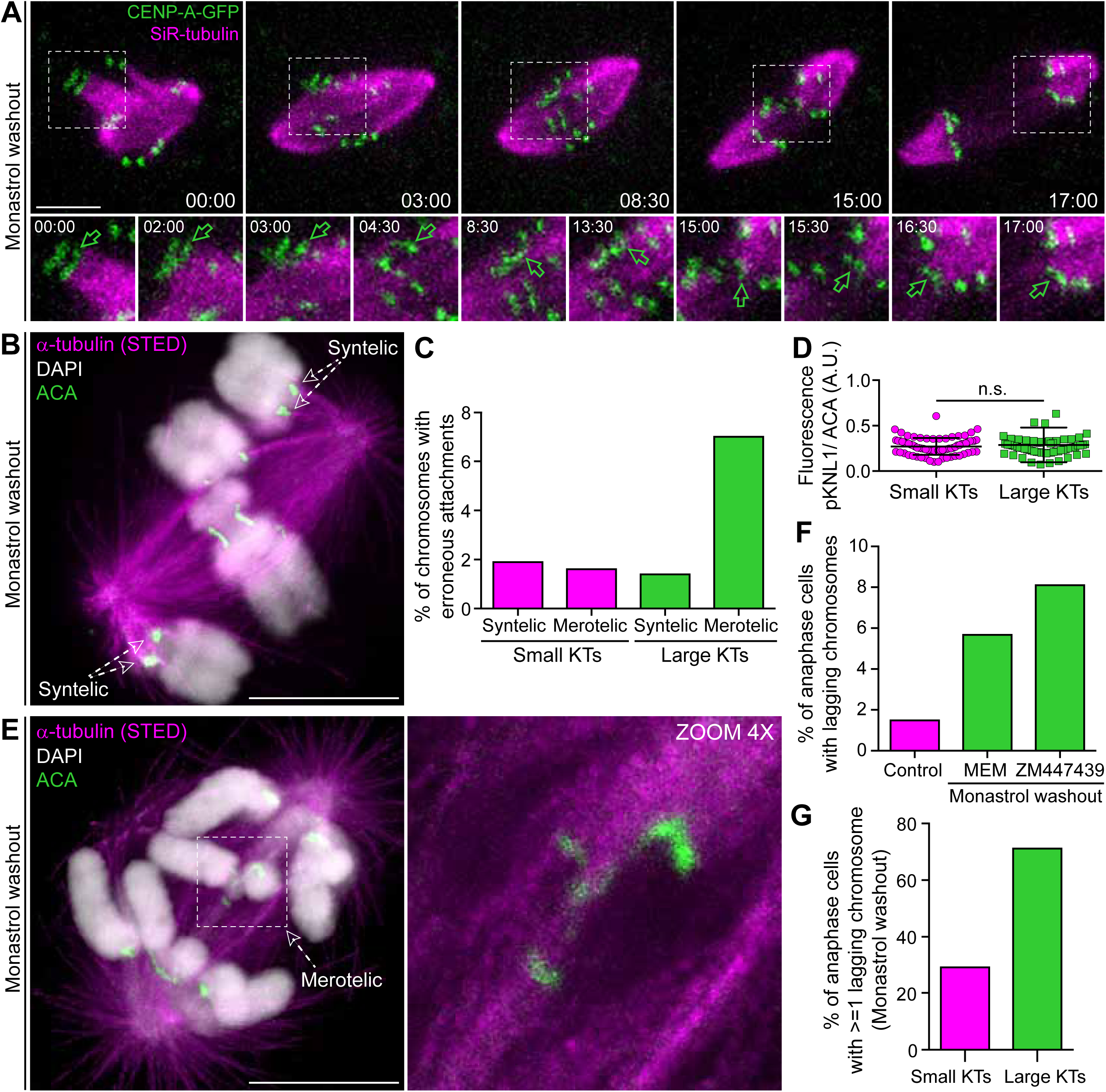
Chromosomes with large kinetochores are more prone to establish erroneous merotelic attachments that result in lagging chromosomes during anaphase. (**A**) High spatial and temporal resolution analysis of error correction after monastrol washout in live Indian muntjac fibroblasts stably expressing CENP-A-GFP (green) and treated with 20 nM SiR-tubulin (magenta). Dashed box highlights a region with a chromosome with a large kinetochore pair (indicated by green arrows in the lower panel). Lower panels show corresponding 1.5x zoom images, plus additional time frames. Note the changes in the conformation of the large kinetochore pair, which lags behind during anaphase, but eventually segregates to the correct daughter. Scale bar = 5 μm. Time = min:sec. (**B**) STED/Confocal image of an IM fibroblast in prometaphase after monastrol washout and immunofluorescence to reveal microtubules (α-tubulin, magenta), chromosomes (DAPI, white) and kinetochores (ACA, green). Dashed arrows indicate one chromosome near each pole with syntelic attachments. (**C**) Quantification of erroneous syntelic and merotelic attachments on chromosomes with small or large kinetochores (KTs) in IM fibroblasts after monastrol washout into Aurora B inhibitor and MG-132. (**D**) Quantification of Aurora B activity at kinetochores of IM as inferred by fluorescence intensity quantification of pKNL1 relative to anti-centromere antibodies (ACA) and normalized for the kinetochore area. There are no statistically significant differences. (**E**) STED/Confocal image of an IM fibroblast in anaphase after monastrol washout and immunofluorescence to reveal microtubules (α-tubulin, magenta), chromosomes (DAPI, white) and kinetochores (ACA, green). A merotelic attachment on a lagging chromatid containing a large kinetochore is indicated (dashed box and arrow). Scale bar = 5 μm. The image on the right shows the corresponding 4x zoom. (**F**) Frequencies of anaphase cells with lagging chromosomes in control IM fibroblasts, monastrol washout into MEM medium and monastrol washout into MEM medium with Aurora B inhibitor. (**G**) Frequencies of anaphase cells with at least 1 lagging chromosome with small or large kinetochores (KTs) after monastrol washout, respectively.

The formation of erroneous merotelic attachments in the early stages of mitosis normally results in lagging chromosomes during anaphase and are a main cause of aneuploidy in mammals [35, 36]. Thus, one would predict that chromosomes with large kinetochores tend to lag more in anaphase when compared with chromosomes with small kinetochores. To test this prediction, and because IM fibroblasts in anaphase showed a very low natural frequency of lagging chromosomes (1.5%; n=267 anaphase cells, pool of 3 independent experiments), we promoted error formation with monastrol as before, and released IM fibroblasts into fresh medium with or without ZM447439 (Fig. 4A, E, F and movie S5). Immunofluorescence analysis in fixed cells after monastrol treatment and washout revealed a significant increase in anaphase cells with lagging chromosomes (5.5%; n=2603 anaphase cells, pool of 9 independent experiments), which was even more pronounced in the presence of ZM447439 (7.7%, n=1323 anaphase cells, pool of 6 independent experiments) (Fig. 4F). Importantly, we found that 73% of the cells after monastrol treatment and washout showed at least one lagging chromosome with a large kinetochore, whereas 30% of the cells showed at least one lagging chromosome with small kinetochores (n=32 cells) (Fig. 4G). Stimulated Emission Depletion (STED) super-resolution microscopy confirmed that the large kinetochores on lagging chromosomes were often found deformed and resulted from merotelic attachments (Fig. 4E). Given that chromosomes with large kinetochores only represent one third of all chromosomes, our measurements indicate a strong bias for chromosomes with large kinetochores to lag behind in anaphase. Because chromosome 3+X in female IM is smaller than chromosome 1 (which has a smaller kinetochore), but larger than chromosome 2 (also with a smaller kinetochore), this works as an internal control to exclude that the measured bias for chromosome 3+X to lag behind in IM was due to a decrease in chromosome size [37]. These data directly demonstrate that chromosomes with large kinetochores are more prone to establish erroneous merotelic attachments that result in lagging chromosomes during anaphase.

Taken together, our results show how an intrinsic kinetochore feature - size - impacts chromosome congression, bi-orientation and error-formation during mitosis. Because chromosomes with large kinetochores bi-orient and congress more efficiently (fig. S2), our work explains why certain species with holocentric kinetochores align their chromosomes in the absence of a CENP-E orthologue [9]. On the other hand, because chromosomes with large kinetochores also establish more errors (fig. S2), one prediction from our work is that chromosomes that use the CENP-E pathway for congression should be less prone to missegregate, explaining why the CENP-E pathway might have emerged during evolution. This would work as a selective pressure to maintain chromosomes with large kinetochores during evolution, suggesting that the errors resulting from incorrect merotelic attachments are unlikely to be propagated. Indeed, our live-cell analysis showed that lagging chromosomes with large kinetochores are normally resolved during anaphase (Fig. 4A and movie S5), consistent with previous findings in other systems [36, 38]. Additionally, any potential loss/gain of an autosome in IM (representing a copy of 1/3 of the genome) might seriously compromise cell viability, in agreement with previous work with male IM fibroblasts reporting that chromosome missegregation and aneuploidy was essentially limited to the smallest Y_2_ chromosome [39]. In humans, the Y chromosome, which has the smallest centromere with significantly less CENP-A compared with any other chromosome [19, 21] also missegregates at elevated frequencies [22], and its loss is the most common somatic alteration in men recently associated with shorter survival and higher risk of cancer [40]. At the other extreme, CENP-A domain expansion and overexpression leading to alterations in kinetochore size, structure and function, have been linked with chromosome missegregation and genomic instability in human cancer cell models [20, 41,42]. Thus, chromosome segregation fidelity might be ensured by a species-specific optimal kinetochore size.

## Acknowledgments

We would like to thank Antόnio Pereira for assistance with STED microscopy. This work was supported by CODECHECK grant from the European Research Council, FLAD Life Science 2020, and the Louis-Jeantet Young Investigator Career Award. D.D. established the Indian muntjac system, performed all the experiments and analyzed the data. A.A. performed the SiR-tubulin titration in the H2B-GFP line. P.A. constructed the joint probability table, performed frequency analysis and 4D tracking of kinetochores. H.M. conceived the project, designed experiments and analyzed the data. D.D. and H.M. wrote the paper. The authors declare no conflict of interests.

## Materials and Methods

### Cell culture

Indian muntjac (IM) female fibroblasts were immortalized with human hTERT and were a kind gift from Jerry W Shay, (University of Texas Southwestern Medical Center, Dallas, Texas) (*31*). IM fibroblasts were grown in Minimum Essential Media (MEM) (Gibco, Life Technologies), supplemented with 20%FBS (Gibco, Life Technologies), 2mM L-Glutamine (Invitrogen) at 37°C in humidified conditions with 5% CO_2_. To collect IM fibroblasts we used Trypsin (Gibco, Life Technologies). Because IM fibroblasts tend to become polyploid in culture, for the purpose of our studies we used only diploid cells, which was controlled by counting the number of kinetochores (2n=6).

### Cell transfection

IM fibroblasts were transfected either with human H2B-GFP (from Geoff Wahl lab, Addgene plasmid # 11680) or pSV-IRESneo3-CENP-A-EGFP (kind gift from Patrick Meraldi, University of Geneva) plasmids using Lipofectamine 2000 (Invitrogen) to generate stable cell lines. For this purpose, at day 1 cells were seeded in 6-well plates at 60-70% confluence in MEM containing 20% FBS. The day after, cells were washed 3x with PBS and incubated with Optimem medium (Gibco, Life Technologies) containing Lipofectamine 2000 (Invitrogen) and the respective DNA for 4h. Optimem with DNA and Lipofectamine 2000 were previously mixed and incubated for 20 min before adding to the cells. After 4 hours Optimem medium was exchanged to MEM supplemented with 20% FBS and cells were selected with G418 (Merck Millipore) after 48h.

### Imunofluorescence

IM fibroblasts were seeded on poly-L-lysine-coated coverslips 2 days before the experiment. After fixation with ice-cold methanol (Invitrogen) or 4% paraformaldehyde (Electron Microscopy Sciences), washed with PBS-0.05%Tween 20 (Sigma-Aldrich) or cytoskeleton buffer pH 6.1 (274 mM NaCl, 10mM KCl, 2.2 mM Na2HPO4, 0.8mM KH2PO4, 4mM EGTA, 4mM MgCl2, 10mM Pipes, 10mM Glucose). Extraction after paraformaldehyde fixation was performed using PBS-0.1%Triton (Sigma-Aldrich). Coverslips were incubated with primary antibodies in blocking solution (10% FBS diluted in PBS-0.05%Tween 20 (Sigma-Aldrich) or in cytoskeleton buffer pH 6.1) for 1h. The following primary antibodies were used: human anti-centromere (CREST) 1:2000 (Fitzgerald), mouse anti-α-tubulin (1:2000; B-512 clone, Sigma-Aldrich), rabbit anti-pKNL1 (1:1000) (*32*) (kind gift from Iain Cheeseman, Whitehead Institute, MIT, Cambridge, MA, USA). Subsequently, cells were washed 3x with PBS-0.05% Tween or cytoskeleton buffer and incubated 45 min with the corresponding secondary antibodies Alexa-488, 568 and 647 (Invitrogen) or Abberior STAR 580 and Abberior STAR 635p (Abberior Instruments) for STED microscopy. For STED microscopy, both primary and secondary antibodies were used at 1:100 concentrations. After adding 1 μg/ml 4´6´-Diamidino-2-phenylindole (DAPI) (Sigma Aldrich) for 5 min, coverslips were washed in PBS and sealed on glass slides using mounting medium (20 mM Tris pH8, 0.5 N-propyl gallate, 90% glycerol).

### Fixed image analysis and acquisition

Image acquisition (0.22 μm thin Z stacks) was performed on a Zeiss AxioObserver Z1 wide-field microscope equipped with a plan-apochromat (1.46 NA 60x) DIC objective and a cooled CCD (Hamamatsu Orca R2). We used Autoquant X (Media Cybernetics) for image blind deconvolution and Fiji (ImageJ) for attachment quantifications and KT tracking. Adobe Photoshop CS4 and Adobe Illustrator (Adobe Systems) were used for histogram adjustments and image preparation for publication. Merotelic attachments were defined by the position of the kinetochores (bent or parallel to the spindle axis) and the respective microtubules (defined through Z stacks). pKNL1 levels were analyzed using ROI manager in Fiji (ImageJ), normalized for the area of each kinetochore. The pKNL1 levels were corrected for the background and normalized to CREST levels (also corrected for the background).

### STED super-resolution microscopy

For STED imaging we used a pulsed gated-STED microscope (Abberior Instruments) with excitation wavelengths at 561 nm and 640 nm doughnut-depleted with a single laser at 775 nm. All acquisitions were performed using a 1.4 NA oil-immersion and a pixel size set to 35 nm.

### Chromosome spreads

IM fibroblasts were incubated with 3.3 μM of nocodazole (Sigma-Aldrich) for 6-7 hours, then trypsinized and centrifuged for 5 min at 1200 rpm. The pellet was resuspended in 500 μL of the supernatant, and a hypotonic solution (medium:water 1:1 and 3.3 μM nocodazole) was added drop by drop until the final volume of 5 mL. The mixture was incubated at 37°C for 20 min. After centrifugation the supernatant was discarded and the cells were fixed with Carnoy (methanol (AppliChem Panreac): acetic acid (Milipore Corporation) - 3:1) solution and kept overnight at -20°C. The following day, the Carnoy fixation was repeated and then the cells were spread drop-by-drop onto the slide. DNA was counterstained with 1 μg/ml DAPI (Sigma-Aldrich) for 10 min and the preparations were mounted on slides using Mounting Media (20 mM Tris pH8, 0.5 N-propyl gallate, 90% glycerol). For chromosome spreads with antibody staining IM fibroblasts were incubated with 3.3 μM nocodazole for 6-7 hours, then trypsinized and centrifuged for 5 min at 1200 rpm. The pellet was resuspended in 500 μL of the hypotonic solution containing sodium citrate (Sigma-Aldrich) and FBS (Sigma-Aldrich) and incubated for 30 min at 37°C. Cells were then placed on glass slides using a cytospin 4 centrifuge (Thermo Scientific). Glass slides containing chromosome spreads were fixed with 4% paraformaldehyde and immunofluorescence was performed as indicated.

### Live-cell imaging

IM fibroblasts stably expressing human CENP-A-GFP or H2B-GFP were plated on fibronectin (Sigma-Aldrich) coated 35 mm glass-bottom dishes (14 mm, No 1.5, MatTek Corporation) 2 days before imaging. Before live-cell imaging, cells were cultured in Leibovitz´s-L15 medium (Gibco, Life Technologies). For tubulin staining, we used 20 nM SiR-tubulin cell-permeable dye (*33*) (Spirochrome) and incubated cells for 6-12 h. Live-cell imaging was performed on a temperature-controlled Nikon TE2000 microscope equipped at the camera port with a modified Yokogawa CSU-X1 spinning-disc head (Solamere Technology), an FW-1000 filter-wheel (ASI) and an iXon+ DU-897 EM-CCD (Andor). The excitation optics is composed of two sapphire lasers at 488 nm and 647 nm (Coherent), which are shuttered by an acousto-optic tunable filter (Goochen&Housego, model R64040-150) and injected into the Yokogawa head via a polarization-maintaining single-mode optical fiber (OZ optics). Sample position is controlled by a motorized SCAN-IM stage (Marzhauser) and a 541.ZSL piezo (Physik Instrumente). The objective was an oil-immersion 60x 1.4 NA Plan-Apo DIC CFI (Nikon, VC series), yielding an overall (including the pinhole-imaging lens) 190 nm/pixel sampling. A 1.5x tube lens (optivar) was also used (126 nm/pixel sampling). Eleven 1 μm separated z-stacks were acquired every 2 min while recording IM fibroblasts stably expressing H2B-GFP. For kinetochore tracking we used IM fibroblasts stably expressing CENP-A-GFP, recorded at 30 sec or 60 sec interval and 0.75 μm separated z-stack. The system was controlled by NIS-Elements via a DAC board (National Instruments, PCI-6733).

### Error formation assay

IM fibroblasts were seeded on poly-L-lysine-coated (Sigma-Aldrich) coverslips 2 days before the experiment. Initially, cells were incubated for 12h with 100 μM monastrol (Tocris bioscience) and after that washed out into MEM medium or MEM containing 1μM Aurora B inhibitor (ZM447439, Selleckchem.com) or MEM containing 1μM Aurora B inhibitor and 20 μM MG-132, a proteasome inhibitor (Calbiochem), based on previous reports (*34*). When indicated, cells were fixed with ice-cold methanol (Invitrogen) for 4 min at -20 °C or 4% paraformaldehyde (Electron Microscopy Sciences) for immunofluorescence analysis.

### CENP-E inhibition

To inhibit CENP-E, IM fibroblasts were treated with 20 nM GSK923295 (*35*) (Selleckchem.com) 1h before fixation or live-cell imaging.

### Frequency analysis and joint probability tables

Custom-made scripts were developed in MATLAB 8.1 (The MathWorks Inc.) to perform the frequency analysis for the number of chromosomes with small and large kinetochores staying at the pole upon CENP-E inhibition. Joint probability tables were calculated for six independent sets of mitotic cells, each with at least 100 cells. The tables were used to calculate the marginal and the conditional probabilities of the number of chromosomes with small/large kinetochores found at the pole. The random variables considered for the joint probability table were ‘S’, for the number of chromosomes with small kinetochores that stay at the pole, and ‘L’, for the number of chromosomes with large kinetochores that stay at the pole. Binomial distributions were fitted to both random variables, using information about the number of independent components (2 for large kinetochores, and 4 for small kinetochores) and the respective experimental values. All data are represented as the mean ± s.d.

### Kinetochore tracking

Live-cell imaging of IM fibroblasts stably expressing CENP-A-GFP was performed as indicated, every 60 sec, and analyzed after CENP-E inhibition using TrackMate Tool in Fiji (Image J). Initial kinetochore and pole positions at nuclear envelope breakdown were manually tracked in four dimensions (x,y,z,t) using Manual Tracking Tool. Further analyses and plotting were performed using MATLAB to assess the initial position of the chromosomes with large kinetochores relative to the spindle and spindle poles/equator. Data from different cells was pooled together by applying geometric affine transformations (without shear) to generate overlap for the poles location. Initial positions of the chromosomes with large kinetochores were plotted on a standardized geometrical representation of the mitotic spindle.

**Fig. S1.**
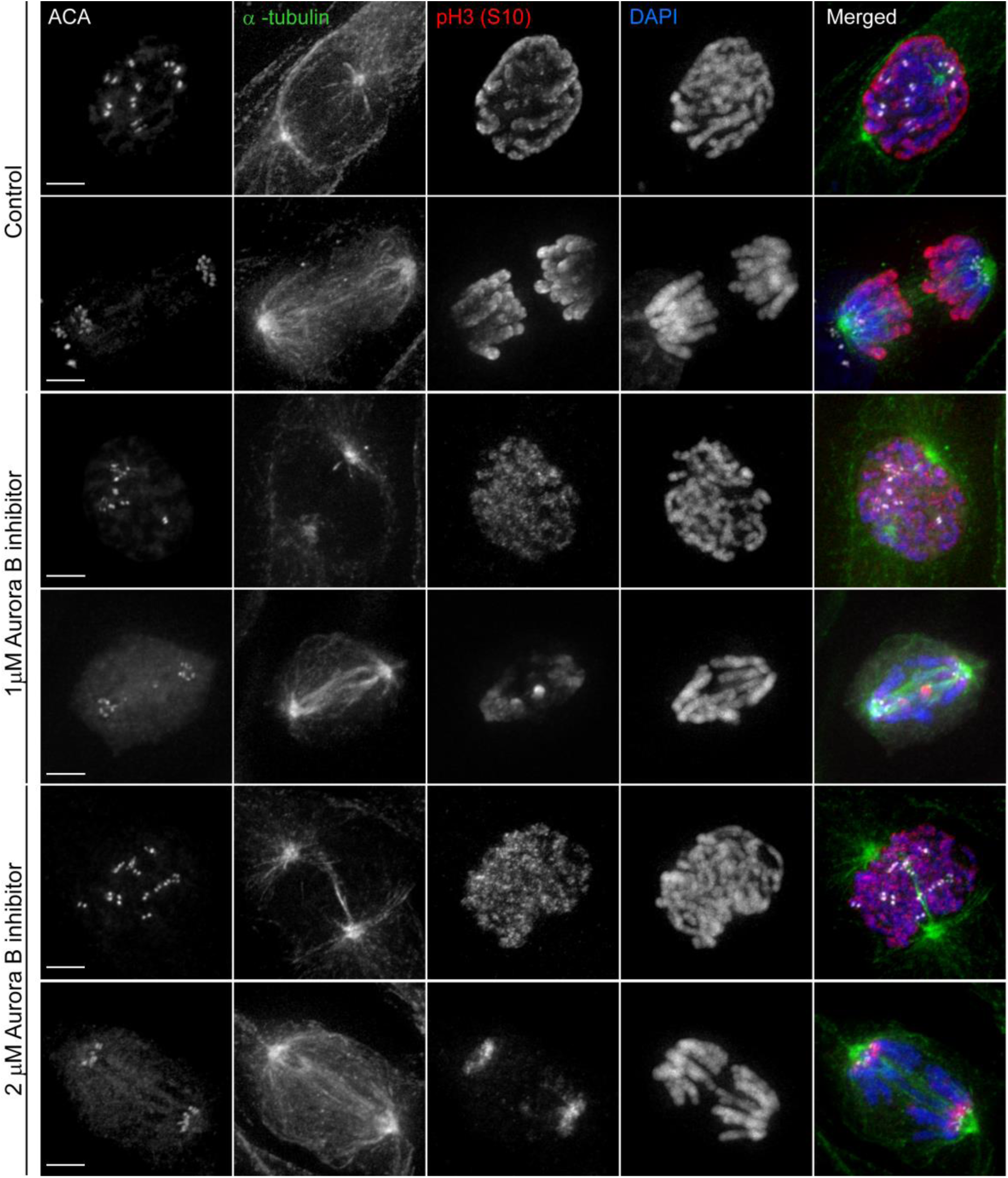
Titration of the Aurora B inhibitor in IM fibroblasts. Immunofluorescence images of IM fibroblasts with or without the Aurora B inhibitor ZM447439 at different concentrations to evaluate the impact on spindle formation and Aurora B activity. In the merged images, kinetochores are shown with ACA (green), Aurora B activity is revealed by pH3(S10) (white), microtubules by α-tubulin (red), and chromosomes with DAPI (blue). Scale bars = 5 μm.

**Fig. S2.**
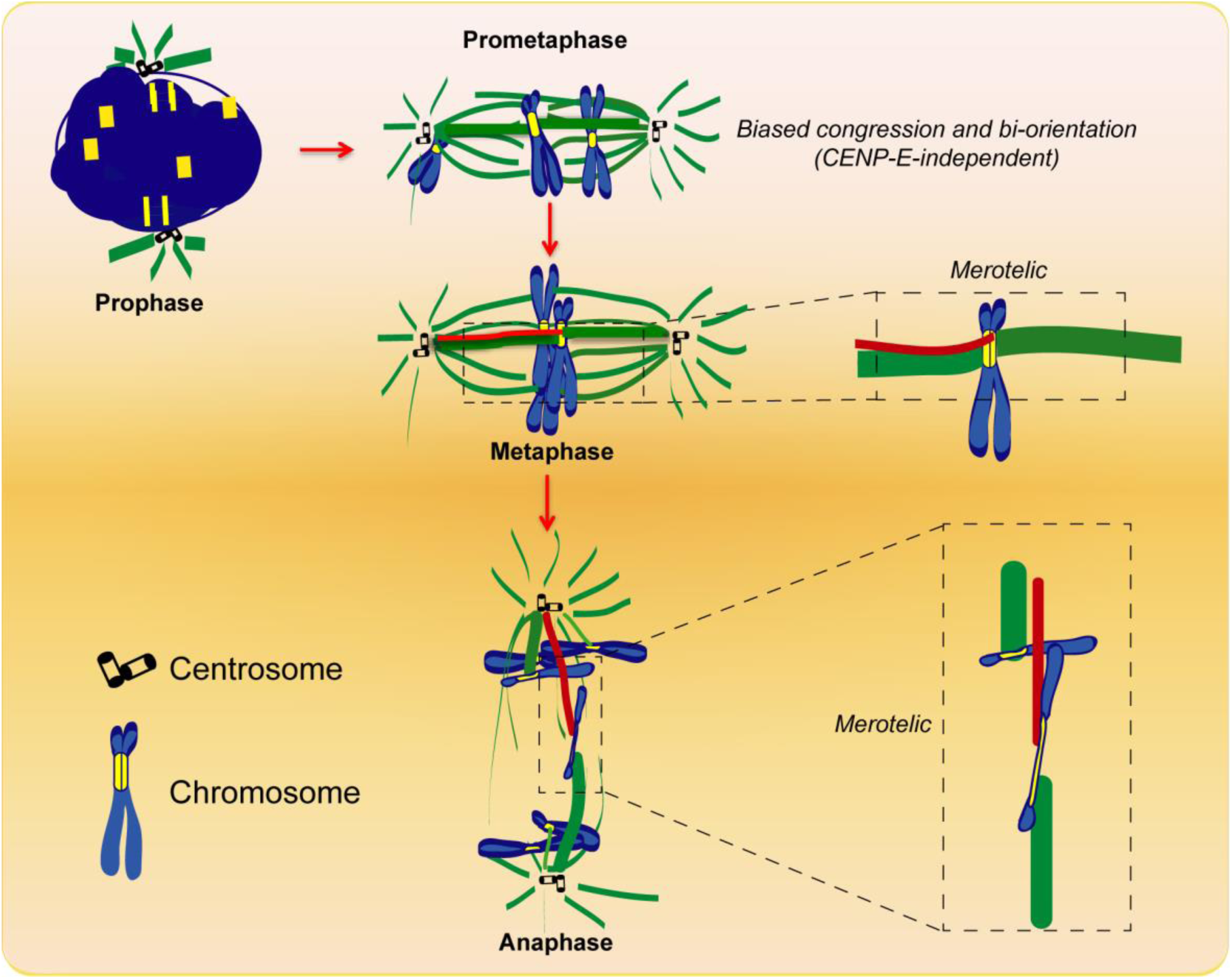
Proposed model for how kinetochore size impacts chromosome congression, bi-orientation and error-formation based on the findings in Indian muntjac. Chromosomes with larger kinetochores would tend to congress and bi-orient independently of CENP-E motor activity. However, they are also more prone to establish erroneous merotelic attachments that would lead to the formation of lagging chromosomes during anaphase and potential missegregation.

**Table S1.** Conditional probability table for the random variables representing the number of chromosomes with small and large kinetochores found at the pole. This table summarizes the experimental data obtained in the CENP-E inhibition experiments. Shaded values correspond to the marginal distributions.

|  |  | # small KTs |  |  |  |  |  |
| --- | --- | --- | --- | --- | --- | --- | --- |
|  |  | 0 | 1 | 2 | 3 | 4 |  |
| # large KTs | 0 | 0.18 | 0.44 | 0.11 | 0.02 | 0.00 | 0.76 |
|  | 1 | 0.10 | 0.08 | 0.04 | 0.00 | 0.00 | 0.22 |
|  | 2 | 0.00 | 0.01 | 0.00 | 0.00 | 0.00 | 0.02 |
|  |  | 0.29 | 0.53 | 0.15 | 0.02 | 0.00 | 1.00 |

## Movie legends

**Movie S1. Mitosis in Indian muntjac fibroblasts.** Live-cell imaging of an hTERT-immortalized IM fibroblast stably expressing H2B-GFP to visualize the chromosomes (green) and treated with 20 nM SiR-tubulin dye to label spindle microtubules (magenta). Time = h:min.

**Movie S2. Tracking individual chromosomes with distinct kinetochore size in Indian muntjac fibroblasts.** Live-cell imaging of a control hTERT-immortalized IM fibroblast stably expressing CENP-A-GFP to visualize the kinetochores (green) and treated with 20 nM SiR-tubulin dye to label spindle microtubules (magenta). Time = h:min.

**Movie S3. Any chromosome can use either the CENP-E-dependent or -independent pathway to congress, regardless of kinetochore size.** Live-cell imaging of hTERT-immortalized IM fibroblasts stably expressing CENP-A-GFP to visualize the kinetochores (green) and treated with 20 nM SiR-tubulin dye to label spindle microtubules (magenta), after CENP-E inhibition with 20 nM GSK923295. Time = h:min.

**Movie S4. Quantitative mapping of the positions of chromosomes with large kinetochores relative to the spindle pole, the metaphase plate and the forming spindle (normalized as an ‘ellipsoid’) obtained by four-dimensional (x, y, z, t) tracking of CENP-E-inhibited cells.** Note that most kinetochore pairs (all green dots) occupy a position at the spindle periphery, but their distribution relative to the pole/equator (red dot) is random. One exceptional kinetochore pair that fell inside the spindle region is also indicated (black dot).

**Movie S5. High spatial and temporal resolution analysis of error correction in Indian muntjac fibroblasts.** Monastrol treatment and washout in a live hTERT-immortalized Indian muntjac fibroblast stably expressing CENP-A-GFP to visualize the kinetochores (green) and treated with 20 nM SiR-tubulin to label spindle microtubules (magenta). Note the difference between chromosomes with a small or large kinetochore pair. One chromosome with a large kinetochore pair can be seen to change conformation and lag behind during anaphase for a short time, but eventually resolves and segregates to the correct daughter. Time = min:sec.

